# Selectivity and dynamics of VAP family tether exchange across membrane contact sites

**DOI:** 10.64898/2026.09.16.752049

**Authors:** Marlieke L.M. Jongsma, Erik Bos, Lennert Janssen, Jimmy J.L. Akkermans, Cami P.M. Talavera Ormeno, Robert Kim, Rayman T.N. Tjokrodirijo, Peter A. van Veelen, Roman Koning, Jacques Neefjes, Ilana Berlin

## Abstract

To integrate diverse cellular function and coordinate homeostasis, the endoplasmic reticulum (ER) communicates with all other intracellular compartments through physical interfaces termed membrane contact sites (MCSs). Given the sheer diversity of possible MCS pairings, how the ER discriminates between simultaneously available binding partners remains unclear. To explore this, we performed comparative endogenous contact site mapping for three closely related VAP family tethers and uncovered a unique set of FFAT motif selectivity and MCS footprints for each: VAPA at mitochondria and TGN, VAPB at mitochondria and peroxisomes, and MOSPD2 at mature endosomes, lysosomes and lipid droplets. In line with their MCS selectivity profiles, co-depletion of VAPA and VAPB, but not MOSPD2, caused systemic breakdown in mitochondrial integrity, while silencing of MOSPD2 alone was sufficient to disrupt the organization and transport of (endo)lysosomes. Unexpectedly, coincident loss of VAPA and VAPB instigated ER contact site rewiring by redirecting endogenous MOSPD2 to the collapsing mitochondrial network. Our findings define hierarchies of ER contact site formation and reveal compensation mechanisms exploited by cells to safeguard organelle homeostasis and crosstalk.

## Introduction

Eukaryotic cells attain enormous biological complexity through compartmentalization of life’s essential processes into functionally distinct membrane-delineated organelles. Extending from the nuclear envelope, the biosynthetic membranes of the endoplasmic reticulum (ER)^1^ feed into the Golgi apparatus^2^, dispensing secretory cargoes to the plasma membrane and other organelles via the *trans*-Golgi network^3^. From here, extracellular materials become engulfed and shuttled through the vesicular network of the endolysosomal system^4^, whose progressive maturation culminates in acidified late endosomes^5,6^ and lysosomes^7^. To fuel these and myriad other cellular activities, mitochondria convert nutrients into energy through oxidative phosphorylation^8,9^, lipid droplets maintain lipid stores^10^, and peroxisomes break down fatty acids and neutralize oxidative stress^11^. To safeguard homeostasis and coordinate effective responses to changing cellular demands, organelles of different identities communicate with and influence one another through regulated membrane appositions (<30 nm)^12^ termed membrane contact sites (MCSs)^13^. Localized exchange of materials and information at MCSs is emerging as a key facet of cellular physiology and resilience across diverse pathways that include Ca^2+^^14^ signaling, lipid^15^ and metabolite^16^ homeostasis, as well as biogenesis^17^ and renewal of organelles^18^, and responses to proteotoxic stress ^19^. Given that different types of MCS pairings arise through closely related mechanisms^20^, whether and how organelles exercise selectivity in MCS formation remains unclear.

The sprawling mesh of the ER membrane network constitutes the largest MCS platform in mammalian cells^21^, housing a wide variety of cross-compartmental tether proteins^22,23^. The best studied in this context is the vesicle-associated membrane protein (VAP) tether family, comprised of VAPA, VAPB, and motile sperm domain-containing proteins (MOSPD) 1-3^24^. All VAPs are integral ER membrane proteins defined by the presence of a cytoplasmic major sperm protein (MSP) domain^25^. VAP proteins exert their membrane tethering functions through MSP-based recognition of FFAT (two phenylalanines in an acidic tract) or FFAT-related motifs found in proteins on opposing organelles. Using proximity-based proteomics of ectopically expressed VAP family proteins, we previously reported that VAPA, VAPB and MOSPD2 share numerous canonical FFAT binding partners^26^. These include (but are not limited to) the Golgi resident tethers ceramide transporter (CERT)^27^ and oxysterol-binding protein (OSBP)^28^; mitochondrial tethers vacuolar protein sorting 13A (VPS13A)^29^ and regulator of microtubule dynamics 3 (RMDN3)/PTPIP51^30^; and (endo)lysosomal tethers OSBP-related protein 1L (ORP1L)^31^ and StAR related lipid transfer protein 3 (STARD3)^32^, among others. Correspondingly, VAPA^33^, VAPB^34^ and MOSPD2^35^ have all been reported to engage in ER MCSs with the same organelles^36^. This begs the question of how multi-functional VAP tethers with overlapping specificities choose their cellular partners, and to what end. Are these choices stochastic or hierarchical? Are they dictated by binding affinities, protein abundances, and/or post-translational modifications? Or do intrinsic differences in partner preference exist?

To address these questions, we exploited CRISPR knock-in technology to fluorescently label endogenous VAP proteins and observe their interactions with other organelles through ER MCSs in the (near)native cellular context. This approach uncovered striking preferences between endogenous VAP family tethers for MCS formation with the Golgi/TGN (VAPA>VAPB), mitochondria (VAPA and VAPB), endolysosomes (MOSPD2>>>VAPB), peroxisomes (VAPB) and lipid droplets (MOSPD2), in living cells. Notably, these partner preferences became masked upon ectopic expression of family members, underscoring the importance of exploring endogenous protein behaviors in their native cellular context. Furthermore, we found that, upon disruption of endogenous ER – mitochondria contacts, cells prioritize compensation over tether selectivity, as evidenced by redirection of MOSPD2 from endosomes and lysosomes to mitochondria, in the absence of VAPA and VAPB. Taken together, our findings present an atlas of endogenous VAP-mediated contact sites with their partner organelles and reveal functional hierarchies between different types of ER MCS in cellular physiology.

## Results

### Endogenous VAP proteins are enriched at distinct ER:organelle contact sites

To study endogenous VAP protein distribution across ER MCSs with different organelles, we introduced GFP coding sequences into the genomic loci of VAPA, VAPB and MOSPD2 in U2OS (osteosacroma) and MelJuSo (melanoma) cell lines using CRISPR/Cas-mediated genome editing technology (Figure 1A). On-target integration of GFP was validated by immunoblotting (Figure 1B) and confirmed by depletion of endogenous GFP fluorescence (GFP-eVAP) upon siRNA-mediated silencing (Figures S1A and S1B). Unlike the ectopically overexpressed GFP-VAP proteins, which distributed throughout the ER, endogenous GFP-VAPs accumulated at discrete sites along the ER membrane (Figures 1C and S1C). To explore whether such enrichments correspond to contact sites, we observed GFP-eVAPs in relation to their partner organelles using fluorescent dyes for mitochondria (Mitotracker), late endosomes and lysosomes (Lysotracker), and lipid droplets (Lipidspot) (Figures 1D and S2A). To visualize peroxisomes, cells were ectopically expressing a peroxisomal targeting sequence (SRL), and TGN was detected following fixation and immunostaining for Golgin97 (Figures 1D and S2A). Assessment of colocalization with subcellular compartments of interest revealed striking differences between endogenous VAPs. Specifically, VAPA was strongly enriched at ER contacts with the TGN and mitochondria, but not at those with endolysosomes, peroxisomes or lipid droplets (Figures 1E and S2B). Endogenous VAPB was also present at ER contacts with mitochondria but showed far less colocalization with the TGN compared to VAPA (Figures 1E and S2B). Instead, VAPB was enriched at contacts with peroxisomes, which attracted neither of the other VAPs (Figures 1E and S2B).

**Figure 1.**
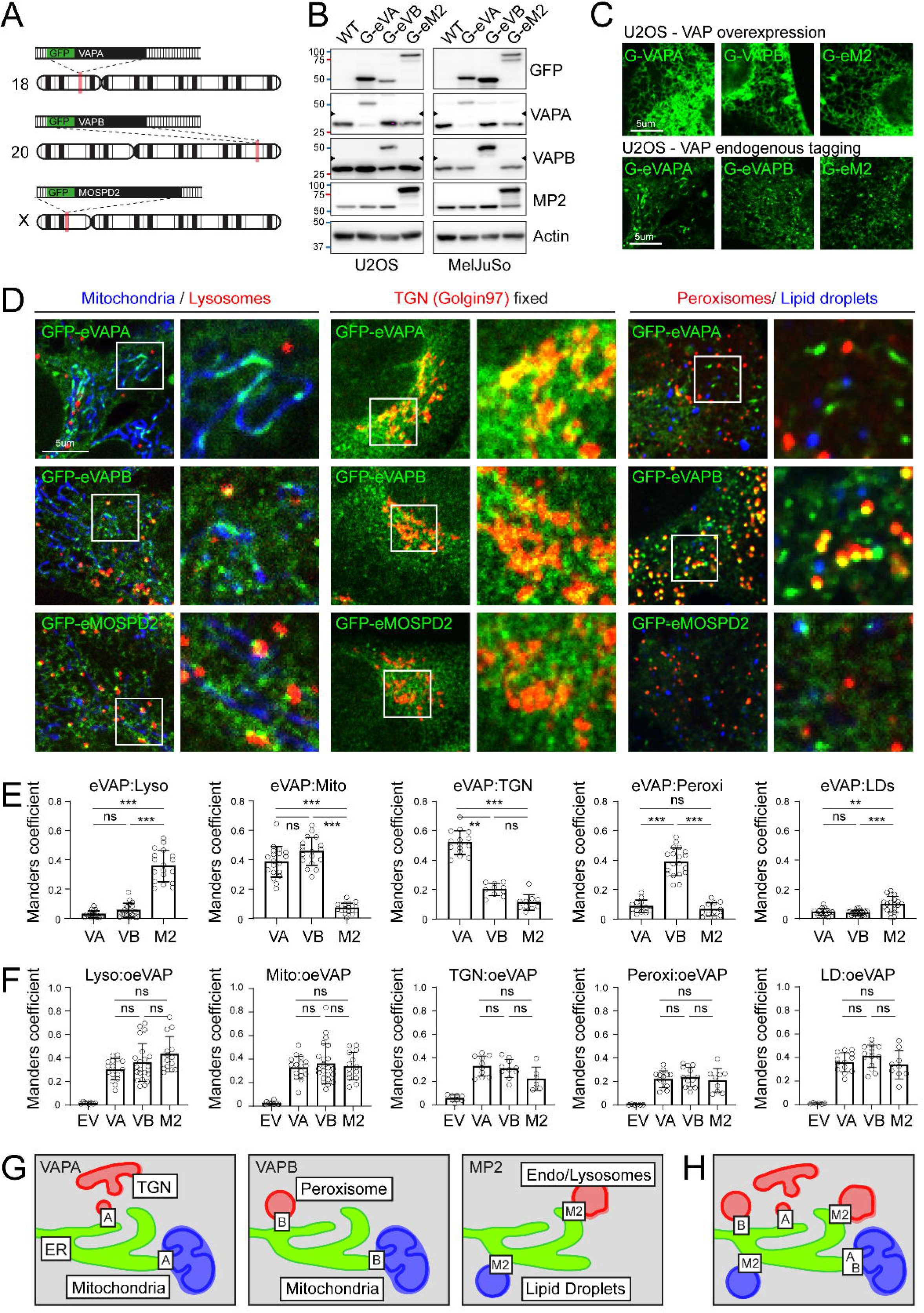
Endogenous VAP proteins are enriched at distinct ER membrane contact sites. **A.** Schematic overview of GFP-tagged VAP genes (GFP-eVAPs). **B.** Immunoblot validation of endogenously GFP-tagged VAP proteins in U2OS and MelJuSo cells. **C.** Confocal images of live U2OS cells overexpressing GFP-VAPs (green, *upper panels*) or endogenous GFP-eVAPs (green, *bottom panels*). Scale bar=5um. **D.** Representative confocal images of U2OS cells endogenously expressing the indicated GFP-eVAPs (green). Stained for (*left panel*) mitochondria (mitotracker, blue) and late endosomes (lysotracker, red), (*middle panel*) immunolabeled against Trans-Golgi-network (golgin97, red), (*right panel*) stained for lipid droplets (lipidspot610, blue) or ectopic overexpression of a peroxisomal targeting sequence (red). Scale bar= 5um. **E, F.** Quantification of colocalization VAPs with different organelles. Graphs report Manders coefficients of endogenous GFP-eVAPs (E) or overexpressed GFP-VAPs (F) with the indicated organellar markers in U2OS cells calculated from single cells, n=3 (TGN n=1-2). **G, H.** Schematic overview of endogenous VAP enrichments at distinct ER:organelle MCSs per VAP (G) or combined (H). *Significance assed using Kruskal-Wallis comparing each column with each column, *** p<0.001, ns: not significant. Error bars reflect +/- SD.* See also Suppl Figures 1-3.

In contrast to VAPA and VAPB, endogenous MOSPD2 was mainly enriched at ER contacts with Lysotracker ^+^ (acidic) late endosomes and lysosomes (Figures 1D, 1E, S2A and S2B). In agreement with a prior report^37^, we observed MOSPD2 as the main VAP present at contacts with LDs, although the prevalence of such contacts was low at steady state, as compared to those with Lysotracker^+^ compartments (Figures 1E and S2B). Contact preferences exhibited by VAPA and VAPB for mitochondria and MOSPD2 for late endosomes/lysosomes were further confirmed in unmodified cells by immunofluorescence using antibodies against different VAPs (Figures S2C and S2D). Moreover, commonalities observed in endogenous contact mapping from two different human cell lines likely reflect generalizable preferences of endogenous VAPs.

Notably, differences between endogenous VAP tethers could not be recapitulated by ectopically overexpressed GFP-VAPs in the same cell lines (Figures 1F and S3A-S3C). These findings underscore the importance of accessing VAP protein localization and behavior at membrane contact sites under endogenous conditions. Taken together, our findings reveal selectivity among VAP family members in communication with different partner organelles (Figures 1G and 1H).

### Favored MSP-FFAT interactions reflect endogenous ER MCS preferences

To demonstrate that the endogenous VAP enrichments observed in proximity of partner organelles reflect *bona fide* ER MCSs, we performed correlative light and electron microscopy (CLEM). Cells endogenously expressing GFP-eVAPA were labeled with Mitotracker and processed for CLEM. Overlays of fluorescence and EM images revealed presence of canonical ER – mitochondria MCS at sites of juxtaposition between GFP and Mitotracker signals (Figure 2A). Next, to explore whether organelle selectivity in contact site formation between endogenous VAPA, VAPB and MOSPD2 could be recapitulated at the molecular level, we isolated GFP-eVAPs from our knock-in cell lines and profiled their co-precipitates by mass spectrometry. Among the top scoring interacting proteins was the mitochondrial ER MCS tether RMDN3, which was preferentially recovered with GFP-eVAPA and GFP-eVAPB as compared to GFP-eMOSPD2 from MelJuSo as well as U2OS cells (Figure 2B and Table S1). Oxysterol-binding proteins primarily localized at the Golgi/TGN (e.g., OSBP and ORP9^38,39^, ORP10 and ORP11^39^) or the plasma membrane (ORP3^40^ and ORP6^41^) followed a similar preference profile (Figure 2B). By contrast, endolysosomal cholesterol transfer OSBL1A/ORP1L was more abundantly recovered with GFP-eMOSPD2 (Figures 2B). Beyond canonical MCS tethers, GFP-eVAPA and VAPB precipitates were broadly enriched in mitochondrial proteins, including the lipid transfer protein STRAD7^42^, a key mitochondrial fission machinery component, dynamin-related protein 1 (DRP1/DNM1L)^43^, mitochondrial fission endopeptidase PITRM1^44^ and ribosomal protein MRPL58/ICT1^45^, among others (Figure 2B). While the endogenous interactomes of VAPA and VAPB exhibited broad overlap in interacting partners form different organelles, some putative associates were able to distinguish between them. Overall, these observations align with the image-based profiling of endogenous VAP:organelle contacts (Figures 1D and 1E) and imply that selectivity in MCS formation is driven by preferential tether pairing.

**Figure 2.**
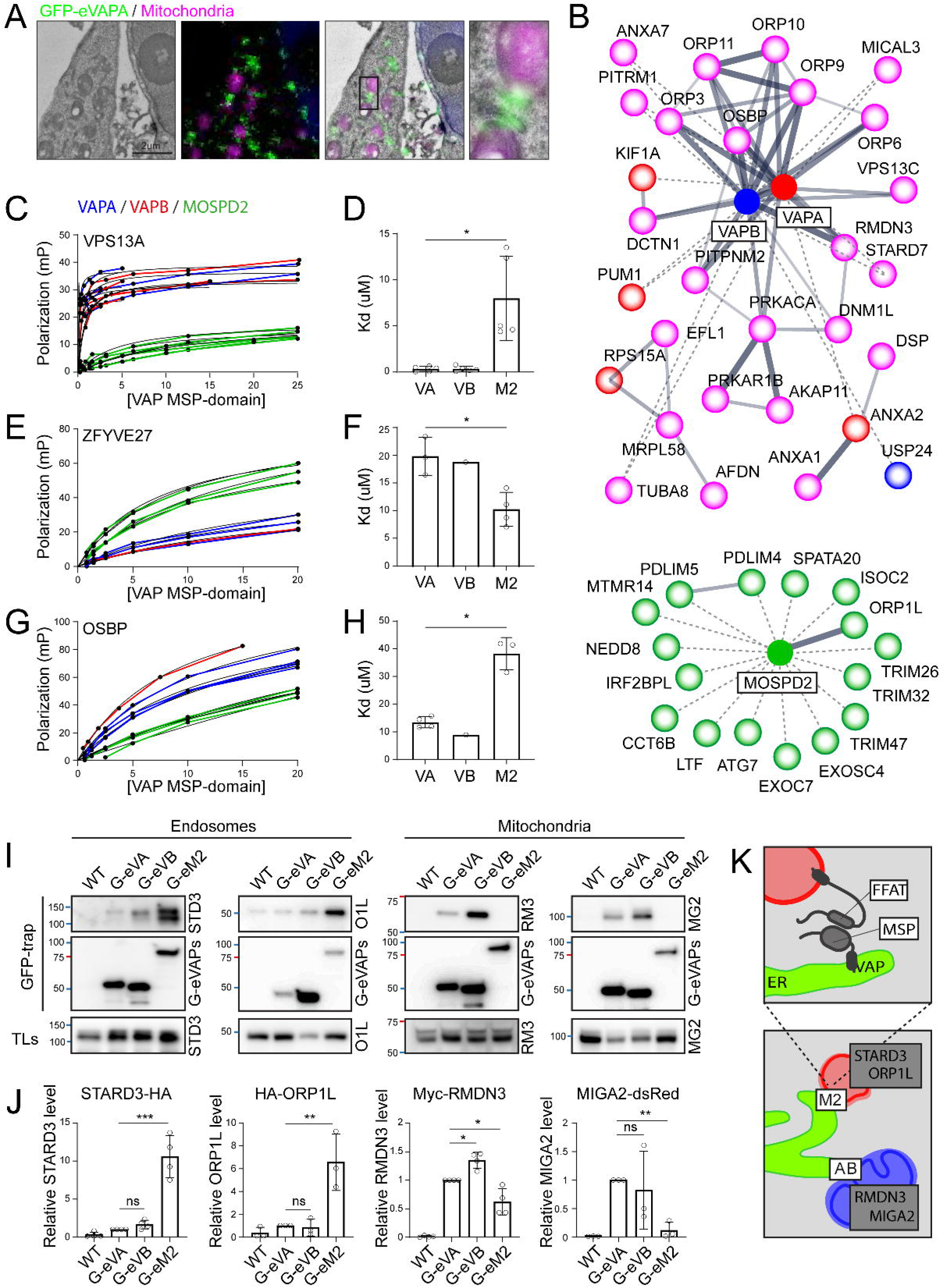
Favored VAP-tether pairings align with endogenous ER MCS preferences. **A.** Visualization of endogenous VAPA:mitochondria MCS using CLEM. Representative overlays of EM and confocal fluorescence images of fixed U2OS cells endogenously expressing GFP-eVAPA (green) and labelled with mitotracker (magenta) are shown. Inset highlights juxtaposition of ER and mitochondrial membranes. Scale bar= 2um. **B.** STRING network analysis of endogenous proteins co-precipitating with GFP-eVAPA (red), GFP-eVAPB (blue) and GFP-eMOSPD2 (green) from MelJuSo and/or U2OS cells. Proteins co-isolating with GFP-eVAPA and GFP-eVAPB are labeled magenta. **C-H.** Fluorescence Polarization (FP) binding assays of VAP MSP domains (VAPA-MSP, blue; VAPB-MSP, red; MOSPD2-MSP, green) to Rhodamine labeled VPS13A (C, D; n=5), ZFYVE27 (E, F; n=1-4) and OSBP (G, H; n=1-4) FFAT-motif containing peptides. Polarization values (mP) were measured as a function of increasing MSP protein concentration. Dissociation constants (Kd) were obtained by nonlinear fitting using a one-site specific binding model. **I, J.** Co-immunoprecipitation of GFP-eVAPs from U2OS and MelJuSo cell lysates with ectopically expressed endolysosomal tether proteins STARD3-HA (n=4) and HA-ORP1L (n=3) versus mitochondrial tether proteins Myc-RMDN3 (n=4) and MIGA2-dsRed (n=3). Representative immunoblots are shown. Quantification (J) reports co-IP relative to GFP-eVAPA. **K.** Schematic summary of preferred interactions between different VAPs and FFAT motif-containing proteins. *Significance assed using Welch’s t test or one-way ANOVA comparing each column to GFP-eVAPA, * p<0.05, **p<0.01, *** p<0.001, ns: not significant. Error bars reflect +/- SD*.

Contact site formation by VAP proteins is typically predicated on MSP domains that recognize proteins harboring FFAT motifs ^24^. To delve further into the mechanism(s) underlying partner organelle selection by individual VAP family members, we compared binding affinities of VAPA, VAPB and MOSPD2 MSP domains for the FFAT motifs of proteins localized on different organelles. Fluorescence polarization assays revealed high affinity interactions between the MSP domains of VAPA (Kd=0.34±0.12 uM) and VAPB (Kd=0.33±0.12 uM) with the FFAT motif of mitochondrial VPS13A (Figures 2C and 2D). The same MSP domains bound with ∼100-fold lower affinities to the FFAT of ZFYVE27/Protrudin (Figures 2E and 2F), a hairpin ER tether known to bridge VAP proteins to the small GTPase Rab7 residing on endosomes and lysosomes^46,47^. In comparison to VAPA and VAPB, MSP domain of MOSPD2 exhibited markedly lower affinity for VPS13A FFAT (Figures 2C and 2D, Kd=8.0±2.0 uM). However, MOSPD2 displayed stronger binding to ZFYVE27 FFAT (Kd=10.3±1.5 uM) relative to its family members (Figures 2E and 2F). These observations fall in line with preferential MCS engagement of mitochondria by VAPA and VAPB and (endo)lysosomes by MOSPD2 (Figures 1D and 1E). Additionally, affinity of MOSPD2 MSP for the Golgi resident OSBP1 FFAT was ∼4-fold lower than that for the FFATs of VPS13A and ZFYVE27 (Figure 2G and 2H), reflecting low contact site formation of MOSPD2 with the Golgi/TGN (Figures 1D and 1E).

Endogenous VAP contactome and MSP-FFAT interaction studies were further validated by co-precipitation of ectopically expressed FFAT-containing tethers with endogenous VAPs. Specifically, mitochondrial RMDN3 as well as Mitoguardin-2 (MIGA2)^48^ were both preferentially recovered with GFP-eVAPA and GFP-eVAPB (Figures 2I and 2J), whereas endolysosomal ORP1L showed better interaction with GFP-eMOSPD2. Although, STARD3 was not detected in our proteomic profiling of endogenous VAP interactors, co-IP experiments demonstrated selectivity for MOSPD2 over VAPA and VAPB, mirroring the behavior of ORP1L (Figures 2I and 2J). Taken together, these observations support a central role for MSP-FFAT interactions as drivers of tether pairing selectivity in contact site formation (Figure 2K).

### Endogenous VAPs present at ER MCSs are highly dynamic

Formation and dissolution of MCSs are central to the maintenance of dynamic cross-compartmental interplay in cellular physiology. To explore endogenous MCS behaviors in living cells, we monitored endogenous GFP-eVAP fluorescence in U2OS and MJS knock-in cell lines. Substantial recovery of endogenous VAPA (70±5%), VAPB (65±2.5%) and MOSPD2 (50%±2.5) was observed within 100 seconds following photobleaching (FRAP) at their respective MCSs (Figure 3A). Dynamic exchange of VAP tethers at organelle contacts suggested that, under overexpression conditions, exogenous VAPs may compete out their less abundant endogenous counterparts (Figure 3B). In line with this, we found that overexpression of mCherry-VAPB reduced the amount of GFP-eVAPA present at contacts with mitochondria, and vice versa (Figures 3C and 3D). By contrast, enrichment of GFP-eMOSPD2 at late endosomes and lysosomes was not affected by overexpression of mCherry-VAPA or mCherry-VAPB (Figures 3C and 3D). Nor did overexpression of mCherry-MOSPD2 affect distribution of GFP-eVAPA and/or GFP-eVAPB to mitochondria (Figures 3C and 3D). These results indicate that VAPs can displace each other based on shared compartment preferences, as illustrated by the exchange of VAPA and VAPB (but not MOSPD2) at ER:mitochondria contact sites when the dose of one VAP is increased.

**Figure 3.**
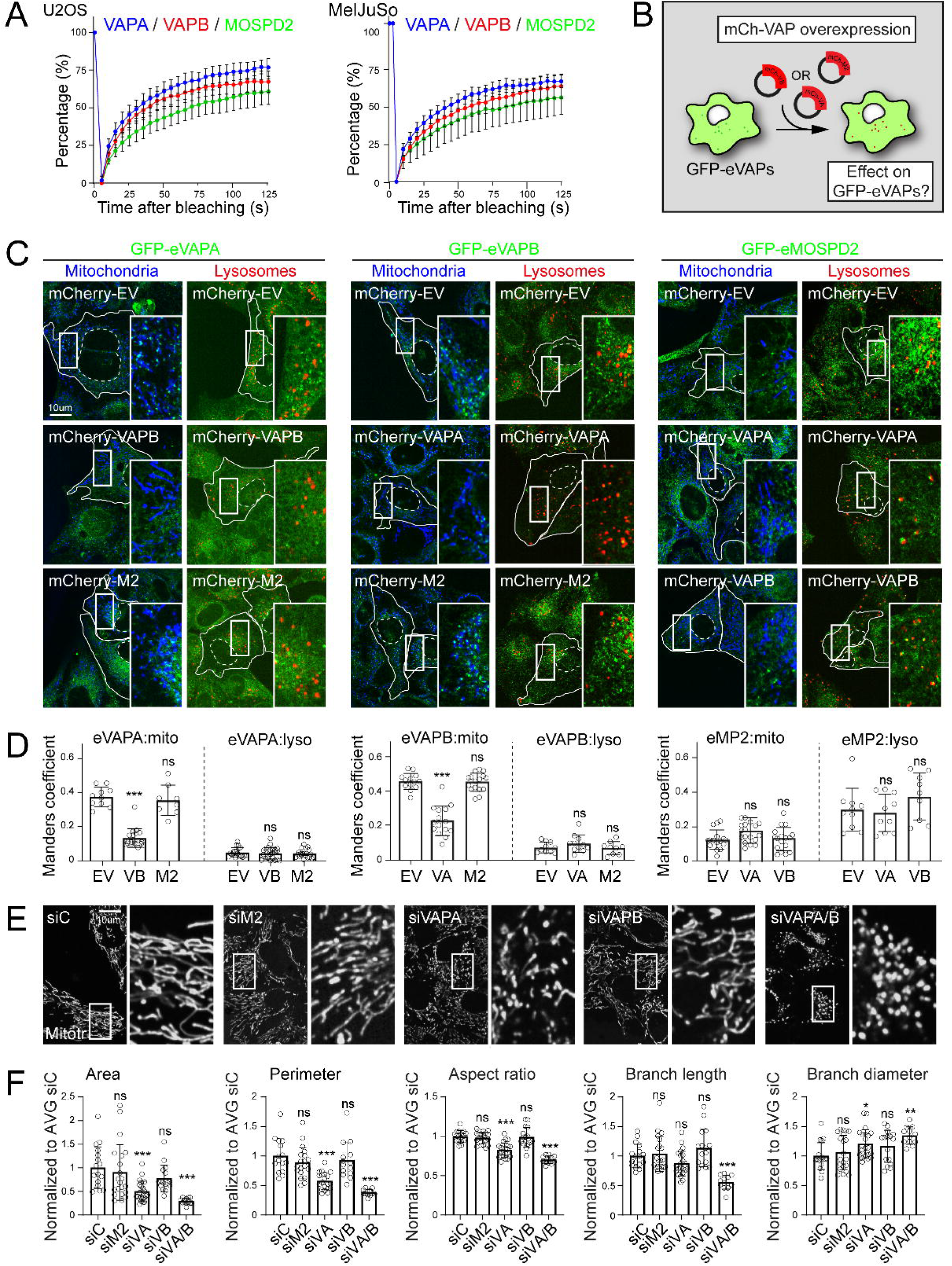
Endogenous VAPs present at ER MCSs are highly dynamic and regulate mitochondrial integrity. **A.** *Fluorescence recovery after photobleaching (FRAP) GFP signal traces for* GFP-eVAPA (blue), GFP-eVAPB (red) and GFP-eMOSPD2 (green)) in U2OS (*left*) and MelJuSo cells (*right*) *collected over 125s post-bleaching are shown. Plotted are average signal intensities from N_VA_=18, N_VB_=20 and N_MP2_=30 cells, n=2.* **B-D.** Effects of ectopic VAP overexpression on endogenous VAPs. **B.** Schematic overview of experimental setup. **C.** Representative confocal images of living U2OS cells endogenously GFP-tagged for indicated VAP (green) ectopically expressing mCherry-empty vector (EV) or mCherry-VAPs (demarcated with solid lines). Stained for mitochondria (mitotracker, blue) and late endosomes (lysotracker, red). Scale bar= 10um. **D.** Co-localization (Manders coefficient) of GFP-eVAPs with mitotracker or lysotracker *as a function of* mCherry-VAP overexpression, n=2. **E, F.** Effects of VAP loss of function on mitochondrial integrity. **E.** Representative confocal images of living U2OS cells transiently transfected with siRNAs targeting the indicated VAPs versus non-targeting control (siC) and stained with mitotracker (gray). Scale bar=10um. **F.** Quantification (MitoAnalyzer, Fiji) of mitochondrial morphology parameters in E. Plotted are average area, average perimeter, median aspect ratio, median branch length and median branch diameter of mitochondria per cell, normalized to siC, n=2. *Significance assed using Kruskal-Wallis comparing each column with each column or one-way ANOVA (compare to siC), * p<0.05, **p<0.01, *** p<0.001, ns: not significant. Error bars reflect +/- SD*.

### VAPA and VAPB regulate mitochondrial integrity

The ER is well established to influence the biogenesis, membrane dynamics and homeostasis of its interacting organelles through MCS formation. We therefore explored the relationship between preferential ER MCS engagement by different VAPs and the organization and dynamics of their partner organelles. To this end, we performed loss-of-function studies using acute siRNA-mediated depletion to avoid long-term compensation effects. Based on VAP displacement experiments described above (Figures 3C and 3D), we hypothesized that VAPA and VAPB share housekeeping responsibilities with respect to mitochondrial architecture and dynamics. In line with this, depletion of VAPA gave rise to aberrations in mitochondrial morphology, and these phenotypes were further exacerbated by combined loss of VAPA and VAPB, but not VAPB only (Figures 3E and S4A). Morphological changes were evidenced by marked reductions in mitochondrial area, perimeter, aspect ratio and branch length, as well as augmentation of branch diameter (Figures 3F and S4B), indicative of mitochondrial fragmentation. Consistent with the absence of contacts between endogenous MOSPD2 enrichments and mitochondria (Figures 1D and 1E), silencing of MOSPD2 had virtually no effect on mitochondrial morphology parameters (Figures 3E and 3F). These observations imply a functional alignment between MCS preferences exhibited by VAP tethers for MCS formation and consequences for the integrity and dynamics of their partner organelles.

### MOSPD2 underpins the organization and dynamics of late endosomes and lysosomes

Distribution and motility of endosomes and lysosomes in cellular space are tightly regulated through physical interactions with the ER^49^. We therefore explored the localization/function relationship between MOSPD2 and the endolysomal system. Endosomes are known to increase physical interactions with the ER membrane along their maturation route^50^. We therefore investigated whether MOSPD2-mediated MCSs reflect a dependence on endosomal maturation. Time-lapse imaging of cells endogenously expressing GFP-eMOSPD2 revealed extensive and stable contact site formation with acidic organelles (Lysotracker^+^) that migrated dynamically in cellular space (Figure 4A). Furthermore, examination of endosomal membrane identity markers in this context demonstrated that vesicles marked by mCherry-Rab7 (late endosome identity) engage in frequent and persistent contact formation with MOSPD2, while those carrying mCherry-Rab5 (early endosome identity) do not (Figures 4B and 4C). Processive contact site engagement during organelle transport has also been illustrated for Protrudin^51^, and may thus reflect a general property of ER – endosome contact sites.

**Figure 4.**
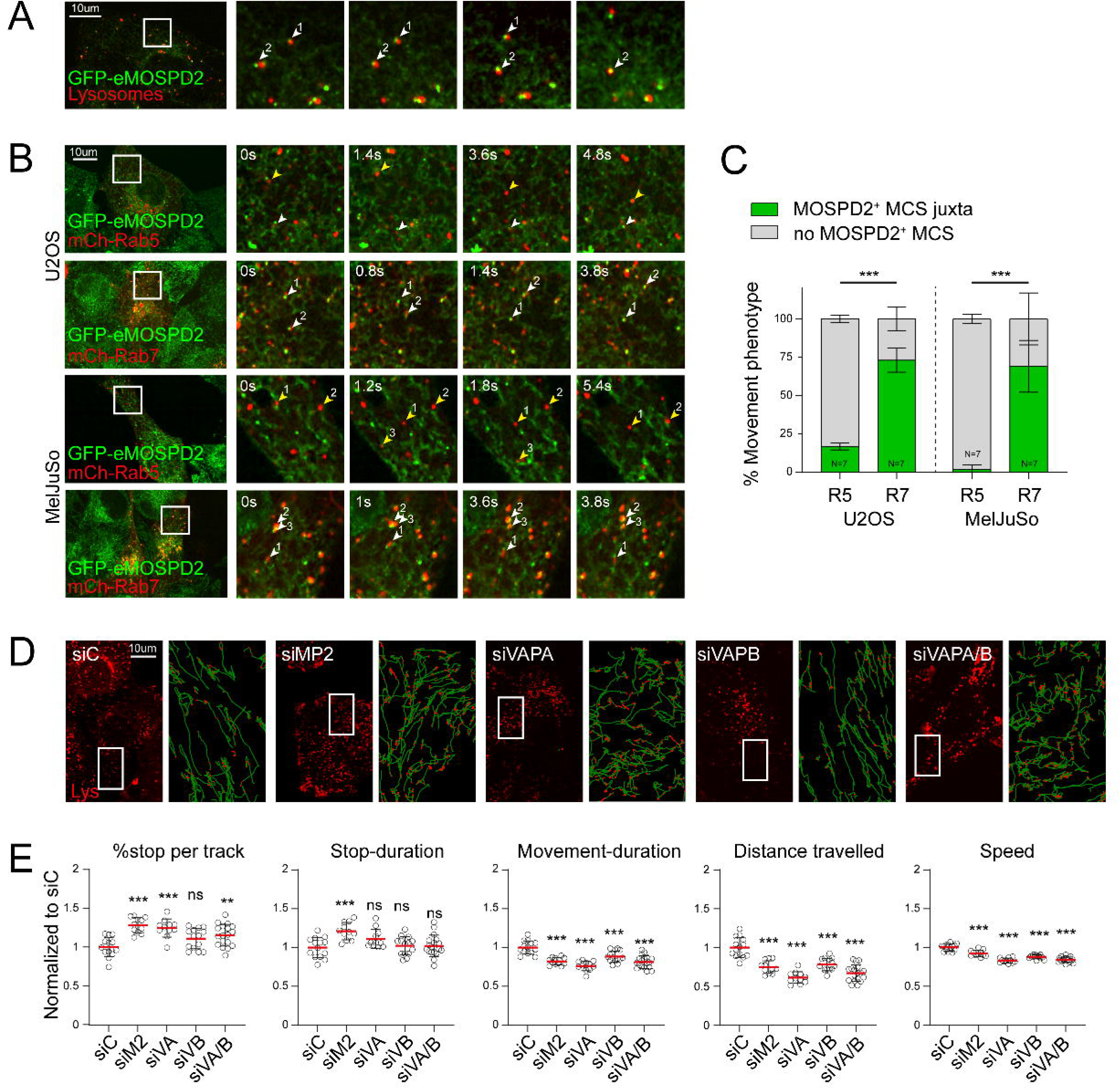
MOSPD2 underpins the organization and dynamics of late endosomes and lysosomes. **A-C.** Time lapse images of U2OS cells endogenously expressing GFP-eMOSPD2 (green) stained with lysotracker (A, red) or transfected with mCherry-Rab5 or mCherry-Rab7 (B, red). White arrows mark ER:LE MCSs; yellow arrows mark endosomes not connected to an ER enrichment. **C.** Quantification of % movement phenotype of endosomes (with or without MOSPD2 enriched MCSs), n=2-3. **D, E.** Effects of VAP loss of function on dynamics of endolysosomes. **D.** Representative confocal images of U2OS cells depleted of the indicated VAPs and treated with lysotracker are shown along with vesicle tracks generated using TrackMate/AnalyseTrackMate. Annotated are movement (green, >0.25um/s) and stops (red, <0.25um/s). Scale bar = 10um. **E.** Quantification of Lysotracker^+^ vesicle track parameters. Plotted are average stop percentage per track, stop-duration, movement-duration, distance travelled and speed of track-fragments per cell, normalized to siC, n=2. *Significance was assed using one-way ANOVA (compare to siC), **p<0.01, *** p<0.001, ns: not significant. Error bars reflect +/- SD*.

To further explore the link between tether specificity and organelle behavior, we proceeded to test the impact of VAP depletion on the organization and dynamics of the endolysosomal system. Under steady state conditions, mature endosomes and lysosomes congregate in a perinuclear cloud ^52,53^ and move bidirectionally to and from the cell periphery in a stop-and-go manner ^51,54^. We therefore investigated the effects of individual VAP family members in this context. Based on our observations casting MOSPD2 as the predominant VAP tether for endosomes and lysosomes, we hypothesized that absence of MOSPD2 would affect the positioning and motility of acidic compartments across cellular space, while loss of VAPA and/or VAPB would have less impact. As expected, silencing of MOSPD2 induced marked dispersion of Lysotracker^+^ organelles throughout the cytosol and resulted in disorganized vesicle transport (Figures 4D, 4E, S4C and S4D). Strikingly, (co-)depletion of VAPA and VAPB also induced endosome dispersion and altered transport dynamics (Figures 4D, 4E, S4C and S4D), despite low prevalence of contacts between GFP-eVAPA and GFP-eVAPB and late endosomes/lysosomes at steady state (Figures 1D and 1E). To reconcile these findings, we tested whether loss of VAPA and/or VAPB alters cellular abundance of MOSPD2. However, no significant changes at the protein level of endogenous MOSPD2 were observed (Figure S4E).

We then hypothesized that loss of individual VAPs could influence endogenous localization of their family members. To determine whether endogenous VAPs respond to loss of their family members, we depleted one VAP at a time and determined the localization of the others. While distribution of GFP-eVAPA to contacts with mitochondria remained unaltered upon depletion of other VAPs, loss of VAPA enhanced mitochondrial recruitment of GFP-eVAPB (Figures 5A, 5B, S5A and S5B). Strikingly, depletion of VAPA caused GFP-eMOSPD2 to partially redistribute toward mitochondria, and this shift was further exacerbated when both VAPA and VAPB were silenced in the same cells, resulting in a dramatic reduction of MOSPD2 contacts with (endo)lysosomes (Figures 5A, 5B, S5A and S5B). Migration of MOSPD2 away from (endo)lysosomes was further evidenced by diminished co-precipitation of STARD3 with GFP-eMOSPD2 and concomitant gain in recovery of mitochondrial RMDN3 and MIGA2 (Figures 5C and 5D). Collectively, these results demonstrate dynamic deployment of endogenous VAPs to available ER docking sites (Figure 5E), implying that cells prioritize ER contacts with mitochondria over those with (endo)lysosomes under conditions when mitochondria-associated VAPs are lost. Such ER contact site rewiring through redirection of MOSPD2 to the collapsing mitochondrial network reveals compensation mechanisms exploited by cells to safeguard organelle homeostasis and crosstalk.

**Figure 5.**
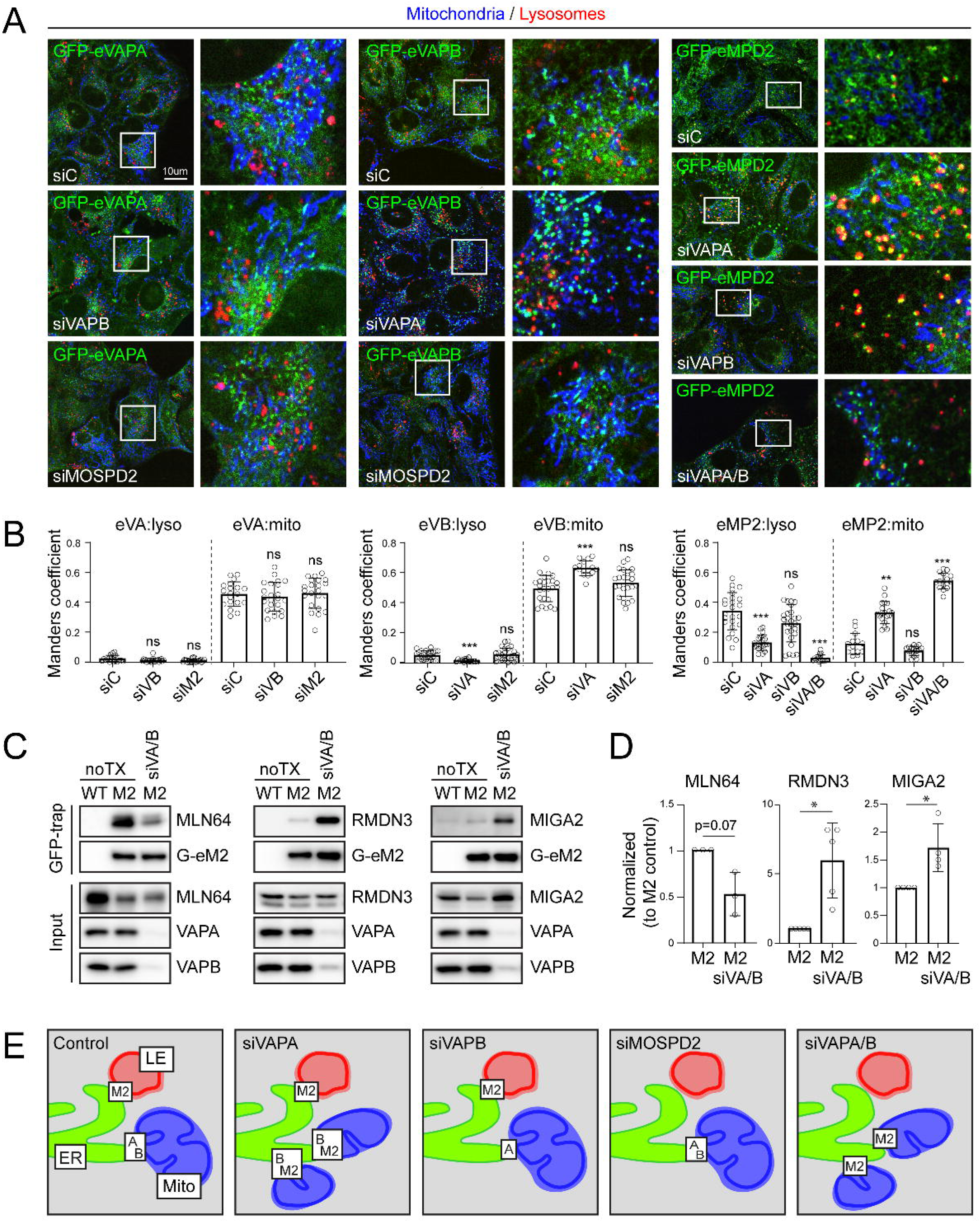
Dynamic VAP MCS rewiring induced by VAPA/VAPB loss of function. **A-E.** Effects of VAP loss of function on family member contactomes. **A.** Representative confocal images and indicated zooms of U2OS cells endogenously expressing GFP-eVAPs (green), depleted for the indicated VAP family members and stained for mitochondria (mitotracker, blue) and late endosomes (lysotracker, red). Scale bar=10um. **B.** Co-localization (Manders coefficient) of GFP-eVAPs with indicated organellar markers in U2OS cells, n=2-3. **C.** Co-immunoprecipitation of GFP-eMOSPD2 from U2OS and MelJuSo cell lysates ectopically expressing STARD3-HA (n=3), Myc-RMDN3 (n=5) or MIGA2-dsRed (n=4) after combined VAPA/VAPB depletion compared to control. Representative immunoblots and depletion controls are shown. **D.** Quantification of data in D. **E.** Schematic overview of endogenous VAP enrichments at ER:LE and ER:Mito MCSs after depletion of other VAPs. *Significance assed using Kruskal-Wallis comparing each column to siC or Welch’s t-test, * p<0.05, **p<0.01, *** p<0.001, ns: not significant. Error bars reflect +/- SD*.

## Discussion

The complex and dynamic patchwork of membrane contact sites is underpinned by direct interactions between different intracellular compartments, each of which features a unique tether repertoire^13^. In this study, we uncovered unexpected diversity among evolutionarily related ER MCS tethers: VAPA, VAPB and MOSPD2. Our findings support a nuanced “selectivity with compensation” model, wherein each tether exhibits a unique footprint in interorganelle communication. At the same time, partial overlap between family members enables homeostatic buffering under conditions of targeted tether suppression. Balancing individual tether preferences against their collective backup capacities thus equips cells with tunable systems-wide organelle integrity controls.

The VAP family of ER tethers has been extensively studied in human physiology and disease^36^. Yet the sheer pervasiveness and interdependence of cross-compartmental interplay in cellular homeostasis have made it challenging to resolve individual tether contributions. Our endogenous contact site mapping reveals that, while both VAPA and VAPB make widespread contacts with mitochondria, neither interacts appreciably with late endosomes or lysosomes at steady state. Instead, together with *Knorr et al.*, we identify MOSPD2 as the principal VAP tether for (endo)lysosomes. Endogenous MOSPD2 also contacts lipid droplets more than the other VAPs, a preference clarified by the unique lipid binding properties of its CRAL-TRIO domain^37^. Meanwhile, endogenous VAPA interacts extensively with the Golgi/TGN, and VAPB shows preferential recruitment to peroxisomes. Considering the functional breadth of ER MCS biology, our findings underscore the importance of accessing endogenous protein localization and behavior at MCS interfaces as a prerequisite for interpreting functional implications of individual VAP family members. In particular, given that nearly all ER MCSs serve as hubs for lipid exchange^55^, and many VAP partners directly participate in lipid transfer between organelles, preferential engagement of different intracellular compartments by endogenous VAPs may dictate local lipid transfer dynamics. Proteomic profiling of endogenous GFP-VAP interactomes and follow-up binding studies broadly support this premise, revealing that lipid transfer proteins residing on mitochondria (e.g., RMDN3^56^ and VPS13A^57^) or the Golgi/TGN (e.g., OSBP^28^ and others) partner more readily with VAPA and/or VAPB, while cholesterol transfer proteins STADR3^58^ and ORP1L^59^, localized on endosomes and lysosomes, preferentially associate with MOSPD2. Though not directly analyzed in the present study, VAP family also mediates ER MCS with the plasma membrane^60,61^, and, in line with published evidence^62^, we identify plasma membrane-localized lipid transfer proteins in precipitates with VAPA and VAPB. Given that the plasma membrane receives the bulk of membrane traffic from the TGN^63^, commonalities in tether use between these compartments would not be surprising. Taken together, these insights position selective tether pairing as the basis for compartmentalization of VAP-mediated MCS engagement at the system’s level, illustrating how three related VAP proteins divide and conquer the complexity of intracellular organelle networks.

Recent advances in imaging capabilities have helped to establish MCSs as highly dynamic liquid-liquid phase-separated structures^64^ supporting rapid tether exchange^65^ of ectopically expressed VAPs. Fluorescence recovery after photobleaching analysis conducted in our genome edited cell lines extends this paradigm by demonstrating that endogenous VAPs are similarly subject to fast exchange. Endogenous VAP enrichments at contacts with other organelles can be envisioned as drivers of nonlinearity in MCS engagement, where multiple medium or even low affinity interactions come together to produce stable binding on the organellar scale. Taking *in vitro* binding data together with genetic perturbations induced to vary cellular concentrations of individual VAPs, we propose that selectivity in MCS formation likely reflects a combined effort of MSP – FFAT binding affinities (individual), enhanced avidity afforded by intramembrane interactions between VAPs (collective), and protein expression of different VAPs (relative). For instance, the MSP domains of VAPA and VAPB bind the FFAT motif of mitochondrial lipid transfer protein VPS13A with sub-micromolar affinities. The same MSP domains show orders of magnitude lower binding to the OSBP FFAT, and even lower affinity for Protrudin, which binds far stronger to MOSPD2. Our observations support a hierarchical model of ER contact site engagement, where different VAPs select their highest affinity binding partners first, thus reducing the available contact pool for the remaining family members. This model explains why VAP overexpression fails to recapitulate endogenous contact preferences, since excess of VAPs bypasses affinity constraints to afford occupancy of all available partners. Hierarchical contact site formation also clarifies the observed shift of MOSPD2 from endolysosomes to mitochondria upon co-depletion of VAPA and VAPB. In this scenario, mitochondrial FFATs (normally saturated by VAPA and VAPB) become available for binding MOSPD2. Conversely, in the endogenous setting, where ER tethers are not in excess of their MCS partners, neither VAPA nor VAPB can rescue the absence of MOSPD2 from endosomes, since higher binding affinities for mitochondrial FFATs (e.g., VPS13A) preclude their migration to lower affinity FFATs on (endo)lysosomes. Collectively, our findings suggest that a simple set of biophysical principles affords sufficient flexibility to balance divergent functions of MCS tethers with (partial) backup capacity.

The ER is extensively documented to regulate the distribution and behavior of other organelles through MCS engagement^23^, exemplified by its ability to decipher the microtubule code and thus control global organelle transport dynamics^66^ and then ride along^67^ to reorganize its own membrane network^68^. Critically, interactions of the ER with mitochondria and endolysosomes dictate fusion and fission of these organelles^69,70^ through spatially and temporally resolved recruitment of membrane deforming machineries at ER MCSs^71,72,73^. Echoing the findings by *Knorr et al.*, we demonstrate that loss of MOSPD2 alone is sufficient to alter endolysosomal system’s architecture and motility, implying a unique function for MOSPD2 in ER-mediated control of proteolytic organelles. Strikingly, combined depletion of VAPA and VAPB (neither of which establish appreciable contacts with endolysosomes under endogenous conditions) proves equally detrimental to the organization and dynamics of these organelles, albeit indirectly. We show that the latter phenotype arises through redirection of endogenous MOSPD2 from endosomes and lysosomes to mitochondria. These findings imply that cells prioritize the function of VAP – mitochondria contact sites over those with proteolytic organelles. Such a choice can be motivated by the catastrophic breakdown in mitochondrial network integrity observed upon coincident loss of primary mitochondrial tethers, VAPA and VAPB. Partial redundancy between VAPA and VAPB in this context parallels that of MFN1 and MFN2^70^ in mitochondrial fusion^74^, with combined perturbations resulting in nonlinear collapse. Given that mitochondrial fragmentation is associated with tissue homeostasis and aging^75–77^, and these findings underscore the importance of VAP-mediated ER – mitochondria contact sites in cellular physiology. In line with broader principles of cellular organization through built-in evolutionary redundancies, our findings shed new light on compensation mechanisms exploited by cells to safeguard essential routes of communication between organelles.

## Materials and methods

### Cell lines

MelJuSo (human melanoma, female) cells were kindly provided by Prof. G. Riethmuller (LMU, Munich) and authenticated by PCR-single-locus-technology (Eurofins genomics, sample code 19 ZE 000486) as MEL-JUSO. MelJuSo were cultured in 7.5 % fetal calf serum (Greiner) supplemented IMDM (Gibco). U2OS (human osteosarcoma, female) cells purchased from ATCC and authenticated by PCR-single-locus-technology (Eurofins genomics, sample code 19 ZE 000484) as U2OS. U2OS were cultured in 7.5% fetal calf serum (Greiner) supplemented DMEM (Gibco). U2OS and MelJuSo cells harboring endogenously tagged proteins (EGFP or mScarlet) were created using CRISPR/Cas9 as described^51^. All cell lines were cultured at 37 degrees, 5% CO_2_ and routinely (negatively) tested for mycoplasma. EGFP-eVAPA (U2OS/MJS), EGFP-eVAPB (U2OS/MJS), EGFP-eMOSPD2 (U2OS/MJS) were generated using the primers, HR and gRNA below.

### GFP-eVAPA

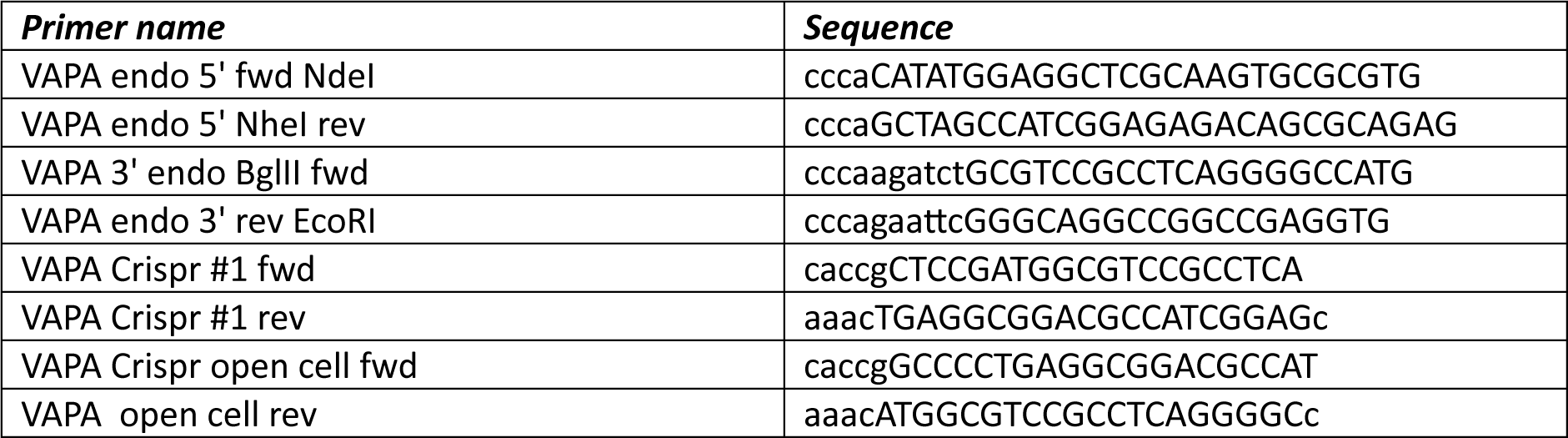

### GFP-eVAPB

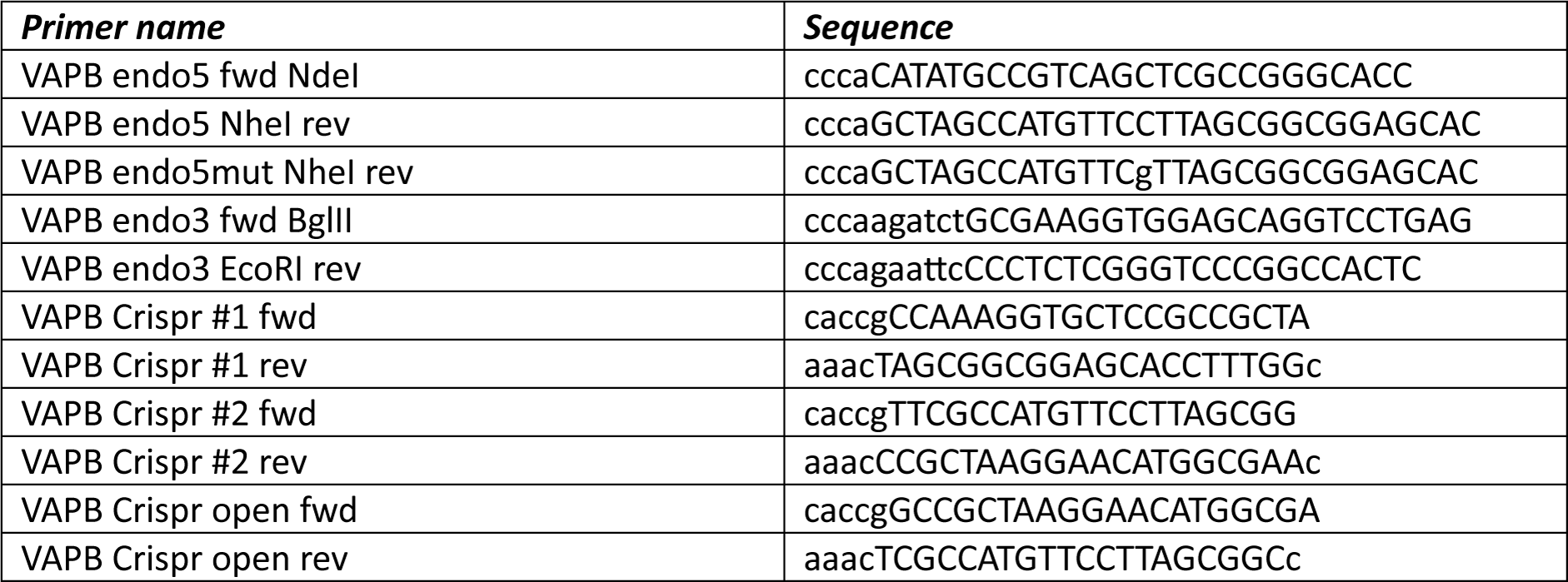

### GFP-eMOSPD2

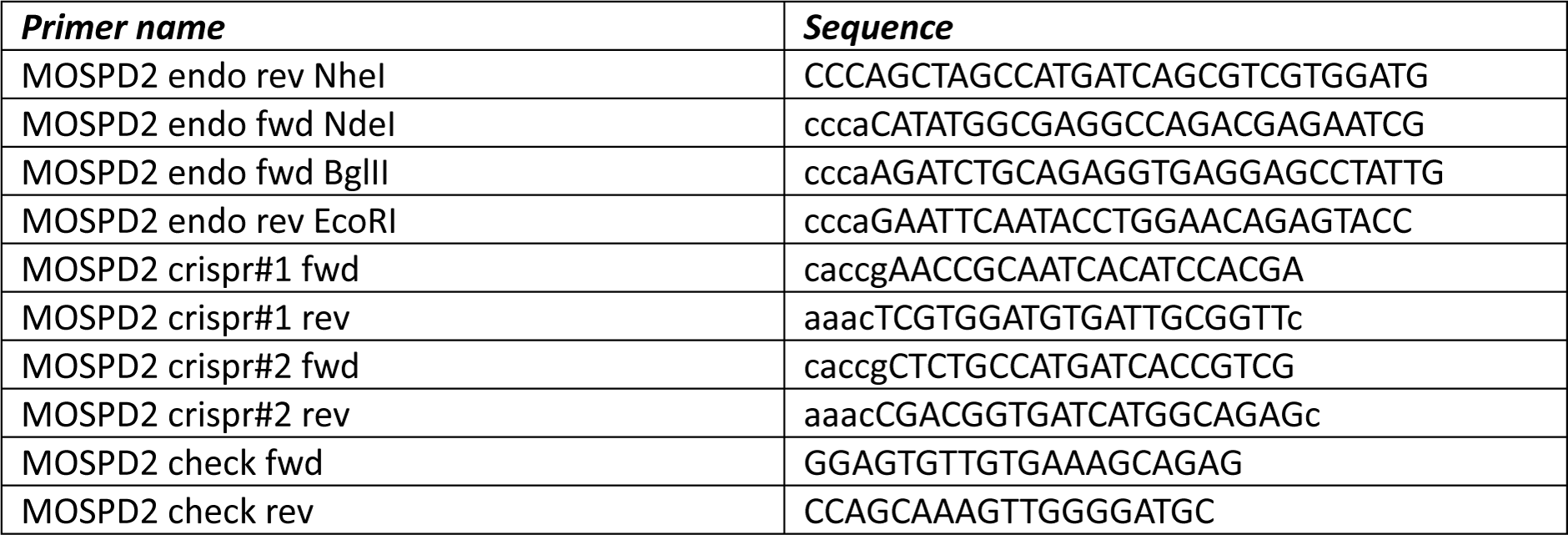

#### Constructs, dyes and antibodies

*(constructs)* pmScarlet-SRL (Addgene #85063), GFP-C1 and RFP-C1 were obtained from Clontech, GFP-VAPA^78^ and GFP-VAPB^78^, GFP MOPSD2^78^ and mCherry-MOSPD2 was obtained by swapping the GFP tag from GFP MOSPD2 for mCherry, mCherry-VAPA (Addgene #226407), mCherry-VAPB (Addgene #108126), STARD3-3xHA (a gift from F. Alpy, IGBMC), HA-ORP1L^79^, n-Myc-RMDN3 (Addgene #227878), hMIGA2-dsRed (Addgene #192866). *(dyes)* MitoTracker DeepRed (Thermo Fisher, M22426), MitoTracker Red CMXRos (Thermo Fisher, M7512), LysoTracker Red DND-99 (Thermo Fisher, L7528), LysoTracker DeepRed (Thermo Fisher, L12492) and Lipidspot610 (Biotium, 70069-T) were used to stain respectively mitochondria, late endosomes/lysosomes and lipid droplets. *(antibodies)* mouse anti-Golgin97 (Invitrogen, A-21270), rabbit anti-VAPA (15257-1-AP), rabbit anti-VAPB (14477-1-AP), rabbit anti-MOPSD2 (83398-1-RR), rabbit anti-GFP (homemade NKI)^31^, mouse anti-RFP (Chromotek, 6G6-100), anti-HA-PO (Roche, clone 3F10, 12013819001), mouse anti-c-Myc (Sigma, clone 9E10, M4439), mouse anti-CD63 (homemade NKI), mouse anti-Tomm20 (Abcam, ab56783) and mouse anti-β-actin (AC-15, A5441) were used to detect the TGN, VAP proteins (VAPA, VAPB or MOSPD2), GFP-, RFP-, HA- or Myc-labelled proteins, late endosomes, mitochondria or actin respectively. For detection by confocal microscopy, primary antibodies were followed by secondary anti-Rabbit or Mouse Alexa-dye coupled antibodies (Invitrogen). To detect endogenous or overexpressed protein by western blot the primary antibodies were followed by Goat anti-Mouse or Rabbit IgG (H+L) Cross-Adsorbed Secondary Antibody HRP (Thermo Fisher, G-21040 and G-21234).

#### Microscopy

(*Fixed*) Cells were cultured on glass coverslips, fixed in 3.75% formaldehyde (Klinipath) and permeabilized in 0.1% TritonX100 (Sigma). Blocking (30min) and antibody incubations (1hr) were performed in 0.5% BSA/PBS (Chemcruz) with 2xPBS wash between the primary and secondary antibody stains. Prolong Gold antifade reagent (Invitrogen, P36934) was used for mounting. (*Live*) Cells were cultured on 35mm glass bottom dishes (1.5 coverglass, MatTek). Fixed and live imaging was performed by a Dragonfly spinning disk microscope (Andor/Oxford instruments) equipped with a Sona 4.2B-6 camera (Andor/Oxford instruments) and a climate chamber. Images were acquired using a Hcx PL 63x 1.32 oil objective (Leica). (*FRAP*) Fluorescent Recovery After Photobleaching experiments were performed using an iLAS system (GATACA) in combination with an Andor Dragonfly microscope equipped with a Zyla 4.2+ camera (Andor/Oxford instruments) and a climate chamber for sequential images acquisition at 5s intervals for indicated times. Photobleaching was performed for 1s at 100% laser power. Maximum fluorescence recovery (i.e. fluorescence prior to bleaching) was set at 100% and the first post-bleach intensity at 0%; post-bleach intensities were corrected for background and bleaching effects.

#### CLEM

Cells were fixed in PHEM buffer (60 mM 1,4-Piperazinediethylsulfonic acid (PIPES), 25 mM N-(2-Hydroxyethyl)piperazine-Nʹ-(2-ethanesulfonic acid) (HEPES), 10 mM EGTA, and 2 mM MgC1_2_, pH 6.9) supplemented with 2% formaldehyde and 0.2% glutaraldehyde (EM grade, EMS) for 2 h. Fixed samples were embedded in 12% gelatin (type A, bloom 300; Sigma) and cut into ∼0.5 mm³ cubes using a razor blade. The sample blocks were subsequently infiltrated with 2.3 M sucrose in phosphate buffer for 3 h. Sucrose-infiltrated blocks were mounted on aluminum pins and rapidly frozen by plunging into liquid nitrogen, after which they were stored under liquid nitrogen until further use. For ultrathin sectioning, frozen samples were mounted in a cryo-ultramicrotome (Leica) at 158 K and trimmed to obtain a square block face of approximately 200 × 200 μm. Ultrathin sections (∼150 nm) were cut using a diamond knife (Diatome) in combination with an antistatic device (Leica). Section ribbons were collected from the cryo-chamber using a droplet of 1.15 M sucrose containing 1% methylcellulose. After thawing, sections were transferred onto titanium specimen grids pre-coated with formvar and carbon. Grids were incubated for 30 min on 2% gelatin in phosphate buffer at 37 °C, followed by rinsing on droplets of PBS at room temperature. Sections were stained with DAPI, rinsed again with PBS and distilled water, and subsequently washed with 50% glycerol. Grids were then placed on glass slides, overlaid with a small droplet of 50% glycerol, and covered with a #1.5 coverslip, which was secured using pressure-sensitive tape. Following light microscopy (Dragonfly spinning disk microscope (Andor/Oxford instruments) equipped with a Sona 4.2B-6 camera (Andor/Oxford instruments)) grids were carefully removed from the slides, rinsed in distilled water, and embedded in a mixture of 2% methylcellulose and 0.4% uranyl acetate (pH 4.5), followed by air-drying. Transmission electron microscopy was performed using a Tecnai 20 microscope operated at 120 kV on a OneView CCD camera (Gatan) using binning 2 at magnifications between 2.300 x (13.1 nm/pixel and 28.600 x (1.0 nm/pix). Correlation and overlay of light microscopy and electron microscopy images was performed using MAPS software (version 3.14 Thermo Fisher Scientific) using the nuclei staining for guiding the overlays.

#### DNA transfection

Cells were transfected using Effectene (Qiagen) or Extremegene HP (Roche) transfection Reagents (Qiagen). Effectene transfections for live imaging were performed in a 35mm-dish format using 200ul EC Buffer, 0.7ug DNA, 5.3ul Enhancer and 6.7ul Effectene for 50-70% cell confluency. Extremegene HP transfections were perfomed in a 6cm-dish format for immunoprecipitation experiments using 500ul -/- medium + 5ug DNA + 10ul Extremegene for 50-70% cell confluency. Cells were imaged or lysed 12-24h after transfection.

#### siRNA transfections

siRNA oligos targeting the indicated genes and non-targeting siRNA (siC, D-001206-13) were obtained from Dharmacon (siGENOME SMARTpools, see Table). For confocal imaging experiments, gene silencing was performed in a 35mm-dish format using 200µL siRNA (500nM stock) mixed with 4µL Dharmafect#1 (Dharmacon) diluted in 196µL IMDM (Gibco) and incubated for 20min on a shaker at RT. The mixture was combined with 30.000 MelJuSo cells or 60.000 U2OS cells resuspended in 1.6ml IMDM and cultured for four days prior to further analysis. For immunoprecipitation assays gene silencing was performed in a 6cm-dish format using 400µL siRNA (500nM stock) mixed with 8µL Dharmafect#1 (Dharmacon) diluted in 400µL IMDM (Gibco) and incubated for 20min on a shaker at RT. The mixture was combined with 120.000 MelJuSo cells or 200.000 U2OS cells resuspended in 3.2ml IMDM and cultured for three (MelJuSo) or four (U2OS) days prior to further analysis.

| ***Protein name*** | ***siGenome (Dharmacon)*** |
| --- | --- |
| VAPA | M-021382-01-0010 |
| VAPB | M-017795-00-0010 |
| MOPSD2 | M-017039-01-0010 |

#### Co-Immunoprecipitations

Cells were lysed in 400ul lysis buffer (50mM Tris-HCl pH8.0, 150mM NaCl, 5mM MgCl_2_, 0.8% NP40, 5% glycerol + complete EDTA-free protease inhibitors (Roche)) and rotated for 20min. Supernatants obtained following 20min centrifugation at 12.000g were incubated with 10ul GFP-trap beads (Chromotek) under rotation for 1h at 4 degrees. Beads were washed 4x in lysis buffer and during the last wash remaining lysis buffer was removed using a needle before addition of 1x NuPAGE LDS sample buffer/5% b-mercaptoethanol followed by 5min incubation at 95 degrees. Samples were separated by SDS-PAGE for western blotting.

#### Total cell lysis

Cells 80% confluent in a 6cm dish were harvested, lysed for 20min in 400ul lysis buffer (50mM Tris-HCl pH8.0, 150mM NaCl, 5mM MgCl_2_, 0.8% NP40 + complete EDTA-free protease inhibitors (Roche)) under rotation and centrifuged for 20min at 4 degrees max speed. 1x NuPAGE LDS sample buffer (Invitrogen, NP0007) /5% b-mercaptoethanol was added to the lysates.

#### SDS-PAGE

Co-immunoprecipitation and total lysis samples were size separated on a 10% acrylamide gel followed by transfer at 300mA for 3hrs to a PVDF membrane (Immobilon-P, 0.45µm, Millipore). Membranes were blocked by 5% milk (skim milk powder, LP0031, Oxiod) for 30min before incubation for 1hr with primary antibody diluted in 5% milk/PBS and washed 2x quick and 2x for 5min in 0.1% PBS-Tween20. Secondary antibody diluted in 5% milk/PBS was incubated for 30min followed by 2x quick and 2x 5min washing in 0.1% PBS-Tween20. Antibody signals were detected after ECL incubation and imaged using an Amersham Imager 600 or 680.

#### Mass spectrometry

(*gel bands*) Automated in-gel digestion was performed on a AssayMap Bravo Platform (Agilent Technologies) following an in-house digestion protocol. Gel bands were first washed with 200 µL of both acetonitrile (ACN) and 100 mM ammonium bicarbonate (ABC) pH 8.4., followed by reduction using 200 µL of 10 mM dithiothreitol (DTT) in 10 mM ABC pH 8.4 at 60 °C for 30 min, and alkylation using 200 µL of 55 mM iodoacetamide (IAA) in 10 mM ABC pH 8.4 at RT for 30 min. Digestion was performed overnight at 37 °C using 50 µL of a trypsin solution (12.8 ng/µL in 25 mM ABC pH 8.4, Porcine, Sequencing grade, Modified, Merck). Peptides were extracted twice using 150 µL ACN/water (80/20, v/v) containing 1.0% formic acid (FA). Samples were lyophilized and stored at −20 °C until analysis to LC-MS/MS. (*mass spectrometry*) Peptides were dissolved in water/formic acid (100/0.1 v/v) and analyzed by on-line C18 nanoHPLC MS/MS with a system consisting of an Ultimate3000nano gradient HPLC system (Thermo, Bremen, Germany), and an Exploris480 mass spectrometer (Thermo). Samples were injected onto a cartridge precolumn (300 μm × 5 mm, C18 PepMap, 5 um, 100 A, and eluted via a homemade analytical nano-HPLC column (50 cm × 75 μm; Reprosil-Pur C18-AQ 1.9 um, 120 A (Dr. Maisch, Ammerbuch, Germany). The gradient was run from 2% to 40% solvent B (20/80/0.1 water/acetonitrile/formic acid (FA) v/v) in 60 min at 250 nl/min. The nano-HPLC column was drawn to a tip of ∼10 μm and acted as the electrospray needle of the MS source. The mass spectrometer was operated in data-independent (DIA) MS/MS mode, with a HCD collision energy at 30%. The lock mass of 445.12003 (siloxane) was used. In the master scan the orbitrap resolution was set to 120.000 and the scan range was 350-1100 at a normalized automatic gain control (AGC) of 100. The maximum fill time was set to 20 ms. For targeted MS2 the orbitrap resolution was set to 15,000 with a scan range of 175-2000, at standard AGC. The selected mass windows were approximately 11 Th wide, with approximately 9 points acquired across the peaks. Data were processed using Proteome Discoverer 3.2 (Thermo), using Chimerys with default settings. Carbamidomethyl was set as a fixed modification and methionine oxidation was set as a dynamic modification. Fragment tolerance was set to 20 ppm. FDR was set to 0.01.

#### Recombinant protein production

GST-VAP^MSP-His proteins (VAPA, VAPB and MOSPD2) were expressed in E. coli BL21 following induction with 0.5mM IPTG 16 degrees overnight. Bacterial pellets were resuspended in lysis buffer (50mM Tris-HCl pH7.5, 250mM NaCl, 1mM EDTA, 1mM DTT), sonicated (3min total time), and clarified by centrifugation at 30.000xg for 30min at 4 degrees. GST-tagged proteins were purified using a GST-binding column and eluted with 25mM reduced glutathione (GSH). The GST tag was cleaved using GST-C3 protease, followed by reverse GST affinity purification to remove the free GST and uncleaved protein. Final purification was performed by size-exclusion chromatography on a Superdex 75 column. Protein concentration was determined using a NanoDrop spectrophotometer. Purified proteins were aliquoted, snap-frozen in liquid nitrogen, and stored at −80 degrees.

#### Preparation of peptides using Solid-Phase Peptide Synthesis (SPPS)

SPPS was performed in tip filters using a Syro II synthesizer (Multisyntech GmbH, Witten, Germany) using standard 9-fluorenylmethoxycarbonyl (Fmoc) based solid-phase peptide synthesis. Amino acids were double coupled in fourfold excess to preloaded Fmoc amino acid trityl resin (0.16 mmol/g, Rapp Polymere GmbH) in N-methyl-2-pyrrolidone (NMP) for 25 min on a 2 µmol scale. The resin was swelled by addition of 90 µL NMP for 5 min (x2). Fmoc was removed by treatment with 20% piperidine in NMP thrice for 3, 5 & 5 min, followed by five washing steps using NMP. Amino acids were coupled by applying a fourfold excess in the presence of 2 equivalents benzotriazol-1-yloxytripyrrolidinophosphonium hexafluorophosphate (PyBOP) and 4 equivalents N,N-diisopropylethylamine (DiPEA) in NMP for 25 min, followed by three washing steps with NMP. After completion of all coupling cycles, the resin was washed with Et_2_O and dried under vacuum. Resin was coupled with Rhodamine by incubating 2 µmol resin with 100 µL of NMP containing 4 µmol of Rhodamine, 4 µmol benzotriazol-1-yloxytripyrrolidinophosphonium hexafluorophosphate (PyBOP) and 8 µmol DiPEA for 16h at room temperature. After washing with NMP, dichloromethane (DCM) and diethylether (Et_2_O), the resin was dried under high vacuum.

#### FP-assay

DMSO dissolved Rhodamine-peptides (25nM final concentration) and indicated concentrations of VAP MSP-domains were diluted in FP-buffer (50mM Tris-HCl pH8.0, 150mM NaCl, 0,05% BSA and 0,01% CHAPS). Assay was performed in 20ul total volume in a 384w plate (Corning, #3820). Peptide only conditions were included to distract background signal. FP values were measured using a PHERAstar FSX (BMG-Labtech).

| ***Protein name*** | ***Peptide sequence FFAT-domain*** |
| --- | --- |
| VPS13A | \$DDSEEEFFDAPC |
| ZFYVE27 | \$AEPDEEFKDAIE |
| OSBP | \$EDDENEFFDAPE |

\$ = Rhodamine

\$ = Rhodamine
| ***Protein name*** | ***MSP-domain*** |
| --- | --- |
| VAPA-MSP-6xHis | aa5-158 |
| VAPB-MSP-6xHis | aa1-125 |
| MOSPD2-MSP-6xHis | aa249-453 |

#### Image analysis

Images were analyzed using Fiji/ImageJ^80^: Manders coefficients were calculated using Image J (JACoP plug-in) to quantify co-localization (protein A:protein B = amount of protein A overlapping with protein B). TrackMate^81^ (vesicle diameter = 0.8μm; thresholds and other parameters were chosen as appropriate based on control samples; the same parameters were used across all samples and all independent experiments) and AnalyzeTrackMate was used for vesicle tracking and analysis^51^. Mitochondria Analyzer was used for mitochondria shape-analysis. ROI manager was used for FRAP image analysis.

#### Statistics

Error bars correspond to SD of the mean. Statistical evaluations report on welch’s t test (two groups), one-way ANOVA (three or more groups, parametric) or Kruskal-Wallis (three or more groups, non-parametric), as indicated, with *p < 0.05, **p < 0.01, ***p < 0.001 and ns: not significant. N= number of cells; n= number of independent experiments.

## Author contributions

MJ and IB designed the study. MJ performed and analyzed most of the experiments. LJ and JA designed and carried out genome editing. CTO conducted peptide synthesis and RK performed recombinant protein production. EB and RK conducted CLEM analysis. RT and PV carried out mass spectrometry. IB wrote the manuscript with input from all authors. IB supervised the project and JN secured financial support.

## Acknowledgements

This work was supported by the Spinoza grant (00897590) awarded to JN. IB is a principal investigator of the Gravitation Consortium “FLOW” (024.006.036), funded by OCW.

---

**Suppl Figure 1. Related to** Figure 1**. A.** Confocal images of GFP-eVAPs in U2OS (*lefts panels*) and MelJuSo cells (*right panels*) validated by siRNAs targeting the tagged VAP protein. Scale bar=10um. **B.** Immunoblot analysis for depletion efficiency of VAPA, VAPB and MOSPD2 in U2OS and MelJuSo cells with actin as loading control. **C.** Confocal images of live MelJuSo cells overexpressing GFP-VAPs (green, *left panels*) or endogenously GFP-eVAPs (green, *right panels*). Scale bar=5um.

**Suppl Figure 2. Related to** Figure 1**. A.** Representative confocal images of MelJuSo cells endogenously expressing the indicated GFP-eVAP (green). Stained for (*left panel*) mitochondria (mitotracker, blue) and late endosomes (lysotracker, red), (*middle panel*) immunolabeled against Trans-Golgi-network (golgin97, red), (*right panel*) stained for lipid droplets (lipidspot610, blue) or ectopic overexpression of a peroxisomal targeting sequence (red). Scale bar= 5um. **B.** Graphs report Manders coefficients of endogenous GFP-eVAPs with indicated organellar markers in MelJuSo cells calculated from single cells, n=3 (TGN n=1-2). **C, D.** Immunostaining of wild type U2OS (C) and MelJuSo (D) cells for VAPA, VAPB or MOSPD2 with indicated mitochondrial (Tomm20) or late endosomal (CD63) marker after methanol fixation. Scale bar= 5um. *Significance assed using Kruskal-Wallis comparing each column with each column, * p<0.05, **p<0.01, *** p<0.001, ns: not significant. Error bars reflect +/- SD*.

**Suppl Figure 3. Related to** Figure 1**. A, B.** Representative confocal images U2OS (A) and MelJuSo (B) cells ectopically overexpressing GFP-empty vector (EV) or GFP-VAP proteins (green). Stained for (*left panel*) mitochondria (mitotracker, blue) and late endosomes (lysotracker, red), (*middle panel*) immunolabeled against Trans-Golgi-network (golgin97, red), (*right panel*) stained for lipid droplets (lipidspot610, blue) or ectopic overexpression of a peroxisomal targeting sequence (red). Scale bar= 5um. **C.** Graphs report Manders coefficients of overexpressed GFP-VAPs with indicated organellar markers in MelJuSo cells calculated from single cells, n=2. *Significance assed using Kruskal-Wallis comparing each column with each column, * p<0.05, ns: not significant. Error bars reflect +/- SD*.

**Suppl Figure 4. Related to Figures 3 and 4. A, B.** Effects of VAP loss of function on mitochondrial integrity. **A.** Representative confocal images of living MelJuSo cells transiently transfected with siRNAs targeting the indicated VAPs versus non-targeting control (siC) and stained with mitotracker (gray). Scale bar=10um. **B.** Quantification (MitoAnalyzer, Fiji) of mitochondrial morphology parameters in A. Plotted are average area, average perimeter, median aspect ratio, median branch length and median branch diameter of mitochondria per cell, normalized to siC, n=2. *Significance assed using one-way ANOVA (compare to siC), * p<0.05, *** p<0.001, ns: not significant. Error bars reflect +/- SD.* **C, D.** Effects of VAP loss of function on dynamics of endolysosomes. **C.** Representative confocal images of MelJuSo cells depleted of the indicated VAPs and treated with lysotracker are shown along with vesicle tracks generated using TrackMate/AnalyseTrackMate. Annotated are movement (green, >0.25um/s) and stops (red, <0.25um/s). Scale bar = 10um. **D.** Quantification of Lysotracker^+^ vesicle track parameters. Plotted are average stop percentage per track, stop-duration, movement-duration, distance travelled and speed of track-fragments per cell, normalized to siC, n=2. **E.** Correlative immunoblot analysis of VAPA, VAPB and MOSPD2 depleted U2OS and MelJuSo cells with actin as loading control.

**Suppl Figure 5. Related to Figure 5. A.** Representative confocal images and indicated zooms of MelJuSo cells endogenously expressing GFP-eVAPs (green), depleted for the indicated VAP family members and stained for mitochondria (mitotracker, blue) and late endosomes (lysotracker, red). Scale bar=10um. **B.** Co-localization (Manders coefficient) of GFP-eVAPs with indicated organellar markers in MelJuSo cells, n=3. *Significance assed using one-way ANOVA (compare to siC) or Kruskal-Wallis comparing each column to siC, * p<0.05, **p<0.01, *** p<0.001, ns: not significant. Error bars reflect +/- SD*.

