## Supplementary figures and images for "Selectivity and dynamics of VAP family tether exchange across membrane contact sites"

### Supplemental Figure 1

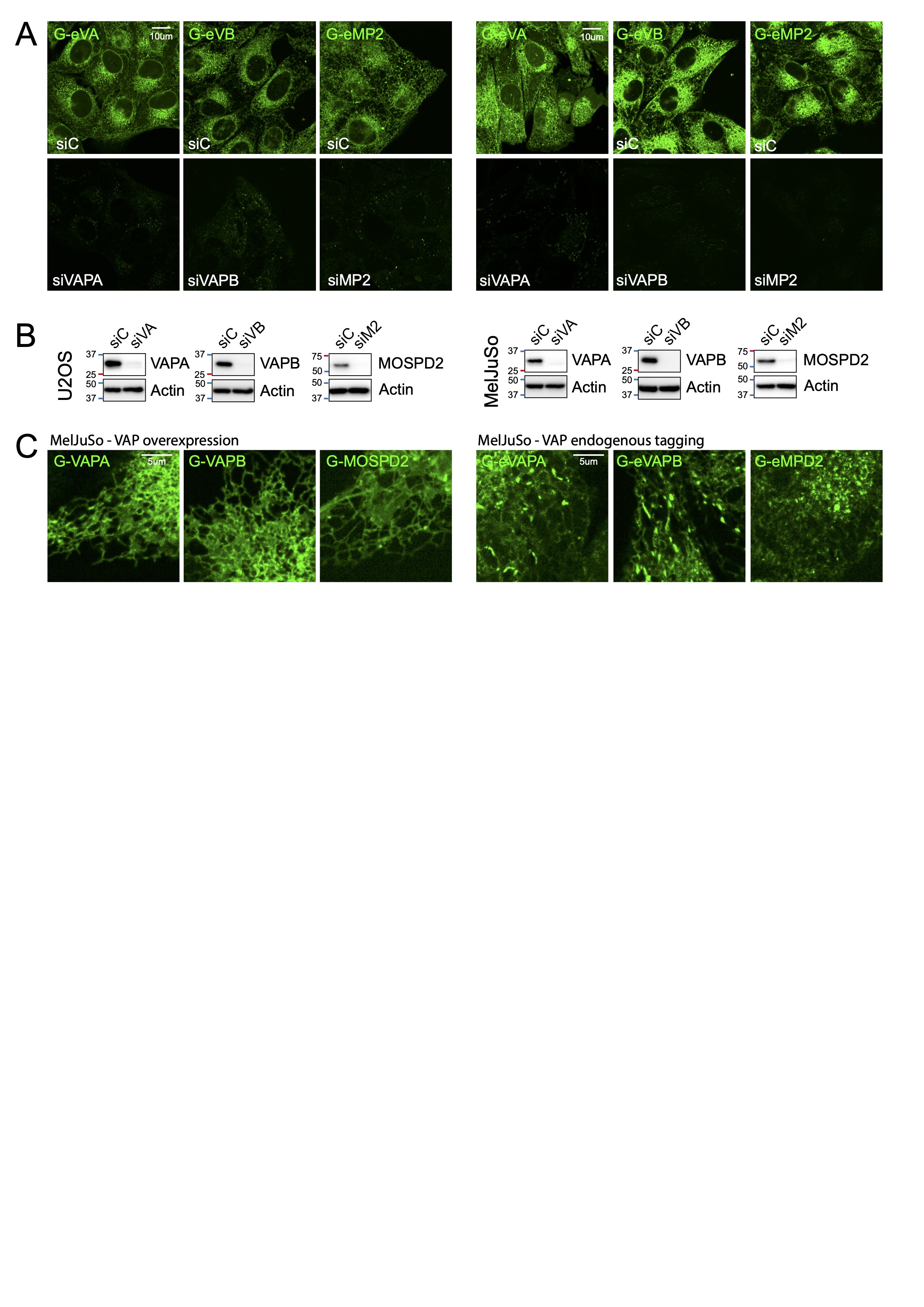

### Supplemental Figure 2

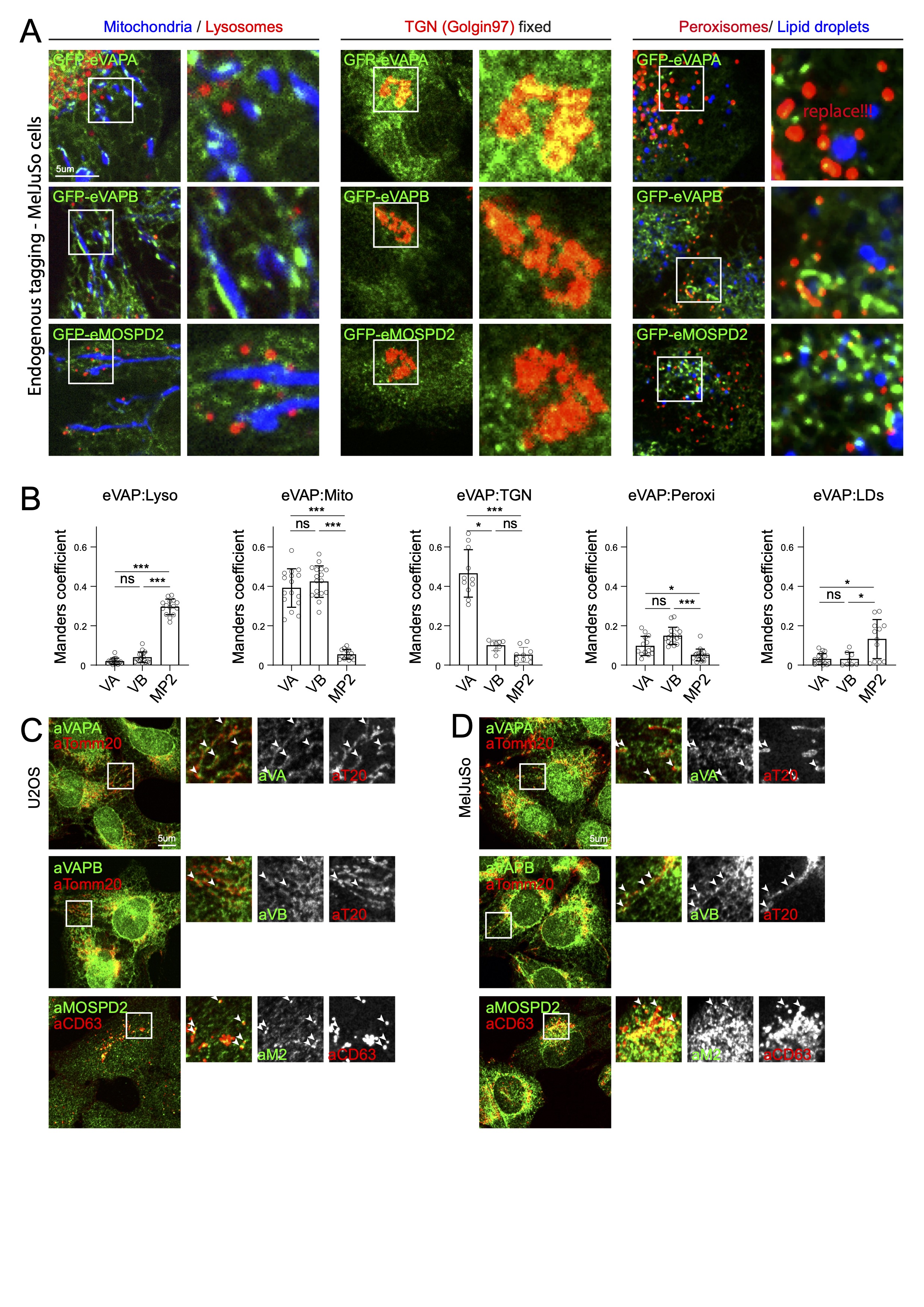

### Supplemental Figure 3

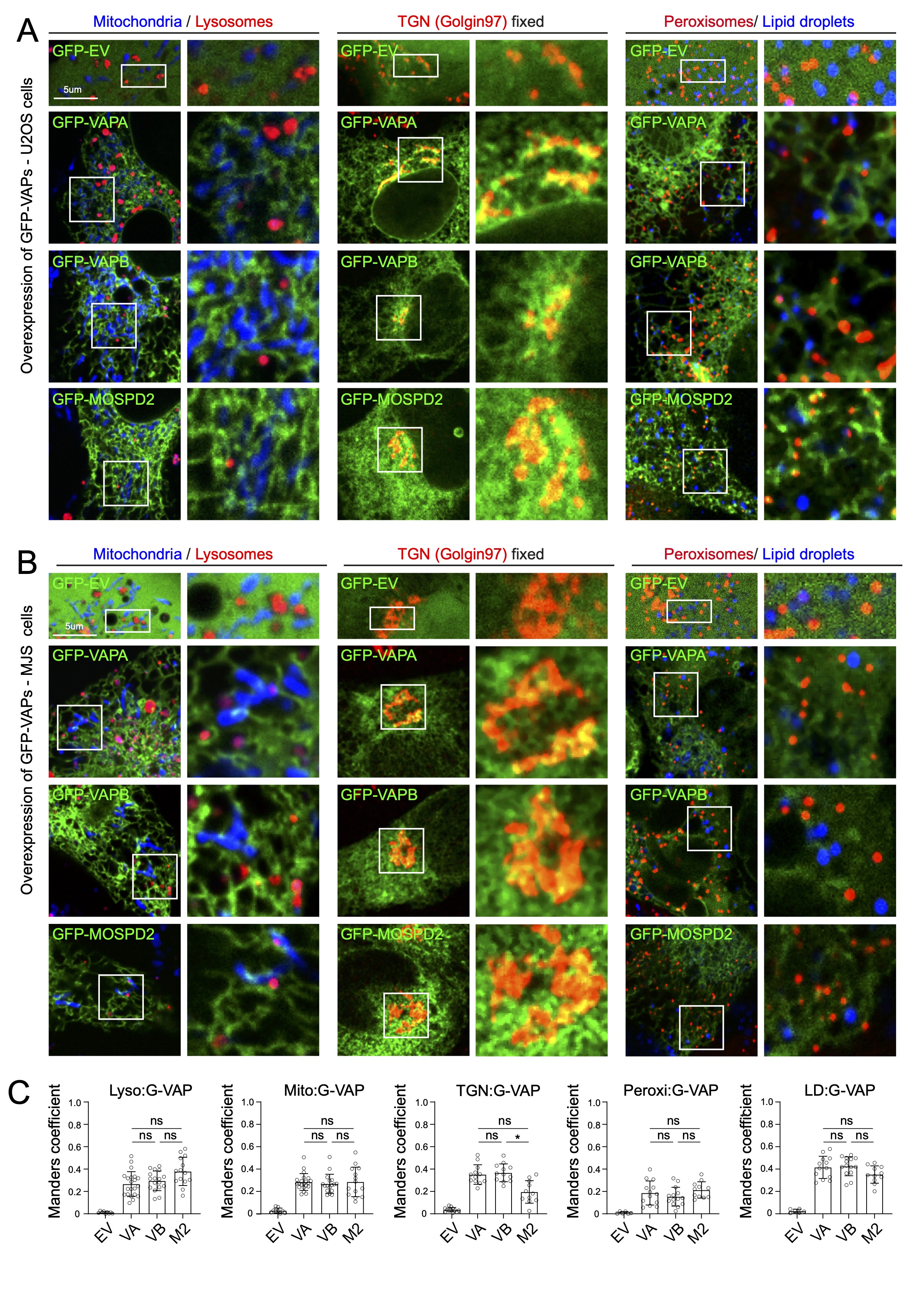

### Supplemental Figure 4

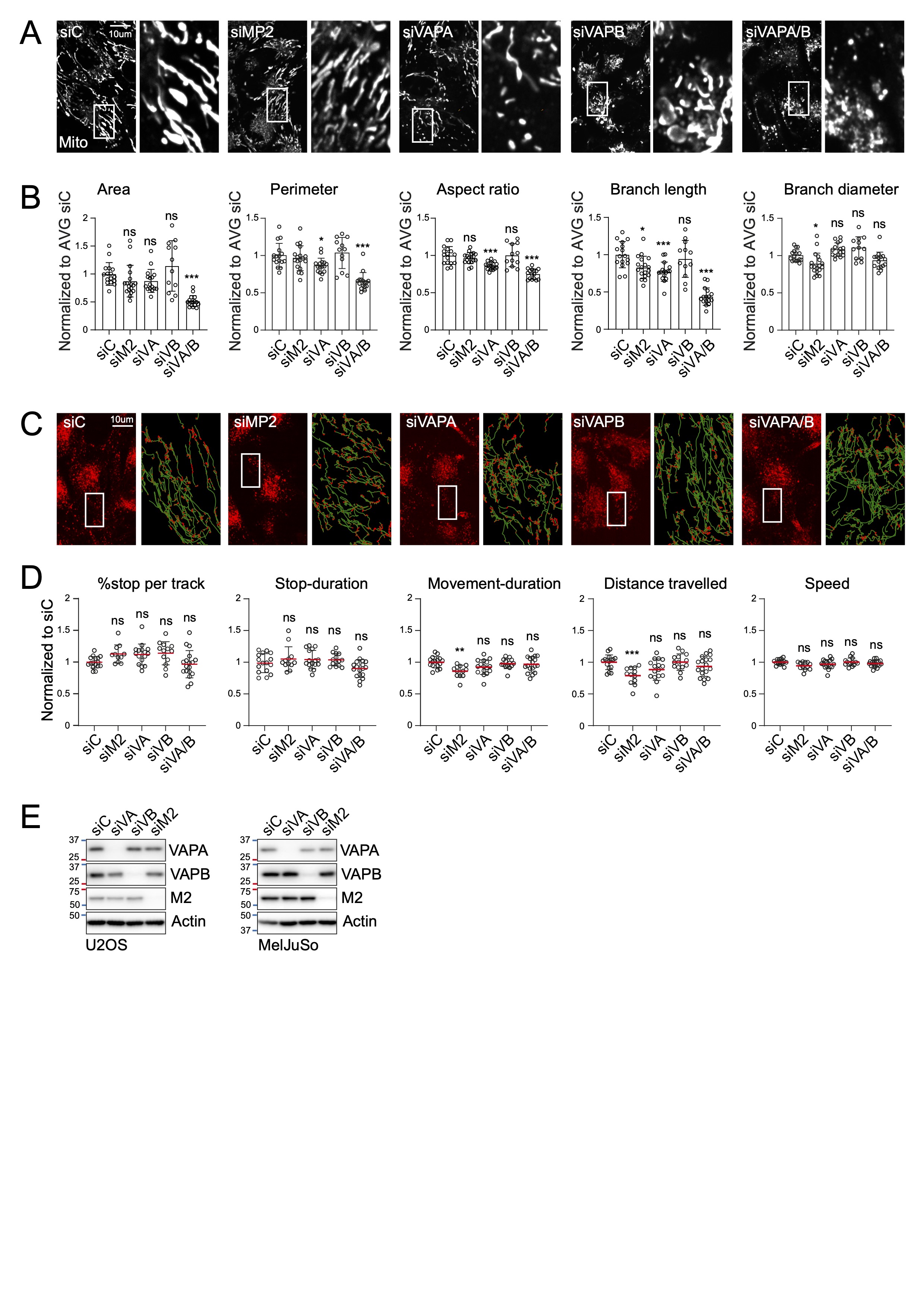

### Supplemental Figure 5

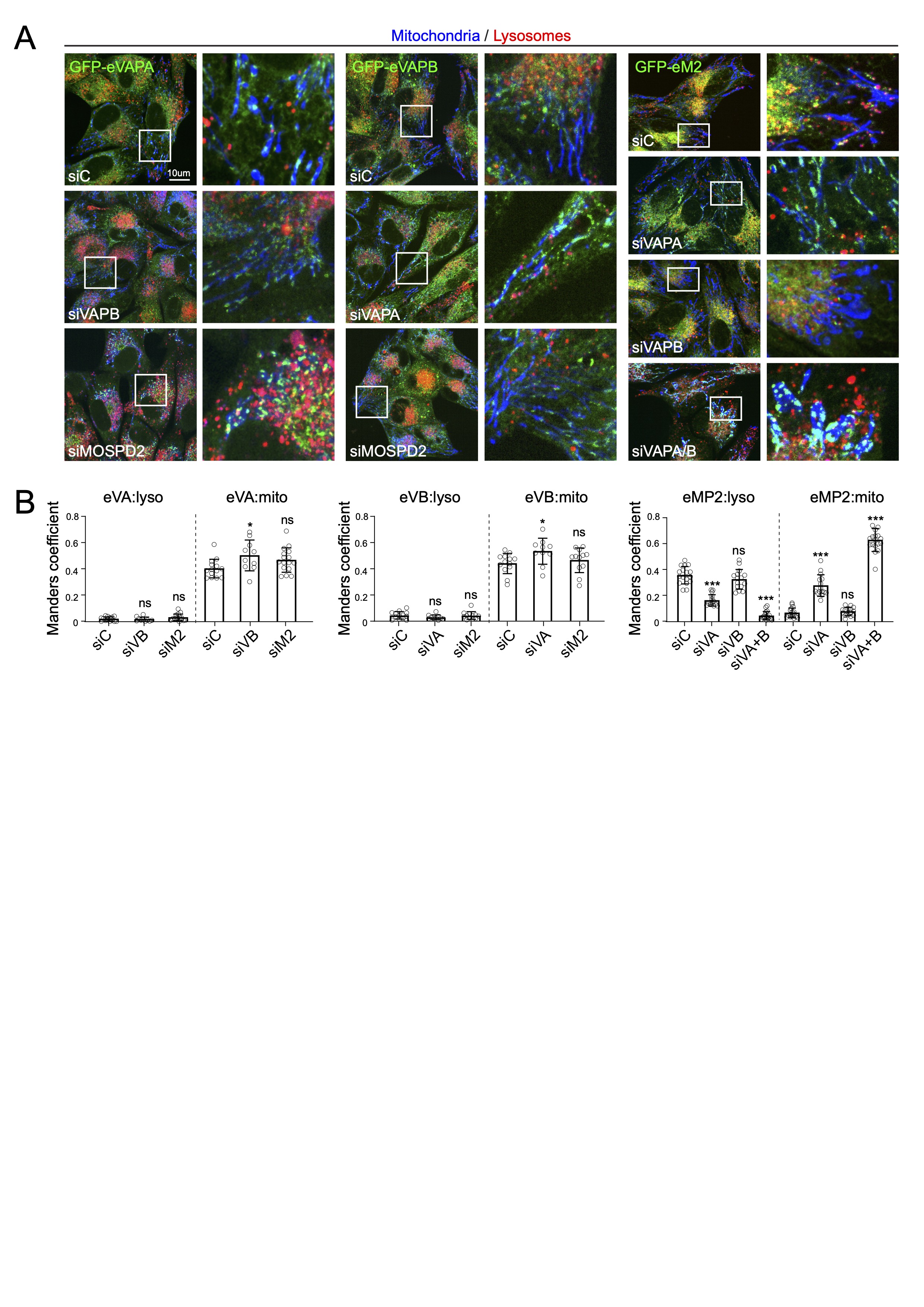
